# Coarse-Grained Modelling of DNA Plectoneme Pinning in the Presence of Base-Pair Mismatches

**DOI:** 10.1101/2019.12.20.885533

**Authors:** Parth Rakesh Desai, Sumitabha Brahmachari, John F. Marko, Siddhartha Das, Keir C. Neuman

## Abstract

Damaged or mismatched DNA bases result in the formation of physical defects in double-stranded DNA. *In vivo*, defects in DNA must be rapidly and efficiently repaired to maintain cellular function and integrity. Defects can also alter the mechanical response of DNA to bending and twisting constraints, both of which are important in defining the mechanics of DNA supercoiling. Here, we use coarse-grained molecular dynamics (MD) simulation and supporting statistical-mechanical theory to study the effect of mismatched base pairs on DNA supercoiling. Our simulations show that plectoneme pinning at the mismatch site is deterministic under conditions of relatively high force (> 2 pN) and high salt concentration (> 0.5 M NaCl). Under physiologically relevant conditions of lower force (0.3 pN) and lower salt concentration (0.2 M NaCl), we find that plectoneme pinning becomes probabilistic and the pinning probability increases with the mismatch size. These findings are in line with experimental observations. The simulation framework, validated with experimental results and supported by the theoretical predictions, provides a way to study the effect of defects on DNA supercoiling and the dynamics of supercoiling in molecular detail.

## INTRODUCTION

The double helical structure of duplex DNA underpins the process of semi-conservative replication while stabilizing the individual strands of DNA and protecting the bases against damage. The helical intertwining of the two DNA strands imposes topological constraints that that must be navigated in all biological processes that involve perturbation of the DNA structure (1). However, these constraints also afford long range control of global DNA conformation via mechanical twisting of DNA (2, 3). The global degree of intertwining of the two DNA strands in the duplex, or equivalently over- or under-winding of the helix is referred to supercoiling(4, 5). *In vivo*, the degree of supercoiling affects cellular processes including gene expression(6), enzyme binding(7) and genome organization(8). Supercoiling is maintained in a state of dynamic homeostasis through the action of a class of enzymes termed topoisomerases, which counteract physiological processes of DNA metabolism such as transcription and replication that alter DNA topology(9).

If dsDNA is topologically constrained, *i.e.* its ends are not allowed to rotate relative to each other, a topological invariant called *linking number* (*Lk*) can be defined: (10, 11)

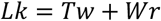

*Lk* refers to the global measure of wrapping of the two DNA strands around each other. This wrapping can be partitioned between twist, *Tw*, the degree of helical intertwining of the two single stands about a common axis, and writhe, *Wr*, the degree of self-wrapping of the dsDNA axis. Torsional stress applied to a piece of linear dsDNA, through a change in *Lk* (by rotating one end relative to the other), results in a change in *Tw* until a critical *Tw*_*critical*_ is reached, beyond which the dsDNA buckles into a structure in which the helical axis of dsDNA wraps or *writhes* around itself forming a “plectoneme”(4, 5).

Supercoiling of DNA is affected by both external environmental factors including the pH (12, 13), tension applied on the DNA(14), concentration of multivalent ions(15); and intrinsic DNA properties such as DNA sequence(16, 17), base-pair mismatches(18), or possibly other structural inhomogeneities. The response of duplex DNA to tension and torsional stress has been extensively studied using single-molecule approaches, principally magnetic and optical tweezers(19). In conjunction with these experimental measurements, theoretical models have been developed to describe the mechanics of torsional DNA buckling and plectoneme formation(20, 21). Computer simulations have also been invaluable for understanding the dynamics and molecular-level details of plectoneme formation(22–29). Despite these advances, many aspects of DNA supercoiling, particularly the effects of DNA defects on DNA supercoiling, remain poorly understood.

Defects in duplex DNA can arise from a variety of factors including base-pair mismatches(18, 30), damaged(31) or modified bases(32), locally melted DNA bubbles(33), or a protein introduced kink in the DNA(34). An elastic rod with a defect that locally decreases the bending rigidity, when subjected to torsional stress, would buckle at the defect(35). However, in genomic DNA that contains millions of base pairs, it is possible that thermal fluctuations and entropic effects could mask the influence of the defect. Alternatively, depending on the type and the size of the defect, the enthalpic gain of buckling at the defect could result in preferential buckling and plectoneme pinning at the defect(18, 36). Furthermore, the type and the size of the defect could significantly alter the structure of the end loop containing the defect; for example, localizing the defect at the sharply kinked tip of a plectoneme may promote the flipping out of the base at the mismatch(37). Indeed, simulations indicate that the distal end of a plectoneme in negatively supercoiled DNA can adopt a sharply kinked conformation in which the base is flipped out in intact DNA(28). Suggestively, some classes of mismatch recognition proteins specifically recognize mismatched or damaged DNA bases through a process in which the DNA is sharply bent and the base is flipped out at the defect site(38).

The importance of a sharply bent DNA in mismatch recognition can also be inferred from a recent simulation study. Sharma et al.(39) quantified the effect of a mismatch on the local rigidity of DNA via all atom simulations. They found that for slightly bent DNA, the presence of a mismatch negligibly affected the bending rigidity. However, when the DNA was strongly bent the decrease in rigidity due to the mismatch became significantly more pronounced. It follows, therefore, that discriminating a mismatch from intact duplex DNA based on energetic differences associated with DNA bending requires a sharp bend. Similar reduction in local bending and torsional rigidity due to the presence of mismatches has also been observed experimentally (40). Given the similarity in the structure of the distal end of a plectoneme observed in simulations and the crystal structure of mismatch recognition proteins bound to mismatched DNA(41–43), it follows that recognition of DNA defects *in vivo* could be enhanced if they were localized at the end of plectonemes due to facilitated buckling. Whereas this is an attractive model to achieve improved rate and efficiency of *in vivo* detection of mismatches, the probability as well as the dynamics of plectoneme formation at mismatches have not been established.

In addition to mismatch-recognition proteins, many other DNA binding proteins introduce a bend in DNA(44), or bind two distal binding sites simultaneously to form a DNA loop, a common motif for transcriptional repression or activation(2). Defects could affect the kinetics and thermodynamics of protein induced bending or looping (45), which could in turn affect transcription and DNA processing more generally. Finally, plectonemes are believed to be able to diffuse along DNA *in vivo* (46) and it is unclear how the presence of mismatches could impact this process.

Here, we study the effect of mismatched base pairs defect on the supercoiling of DNA. Recent single-molecule magnetic tweezers-based experiments by Dittmore *et al.*(18) found that for positively supercoiled (overwound) DNA in the limit of high salt concentration (0.5 M to 1 M NaCl) and high force (>2 pN), a plectoneme forms and remains localized at the location of a single mismatched base pair. This provides a possible mechanism for detection of mismatches by mismatch repair enzymes. However, for technical reasons, the single-molecule experiments could not measure pinning under physiological conditions (0.1 M NaCl and forces on the order of 1 pN or less). Brahmachari et al.(36) developed a statistical-mechanical model describing the localization of plectonemes under physiologically relevant conditions (0.1 M NaCl and 1 pN). The results of these calculations indicate that the plectoneme localization at the mismatch becomes probabilistic under physiological conditions due largely to entropic effects. This model used a simple analytical approach in which the mismatch was assumed to result in a local reduction in the bending energy. This combination of theoretical and experimental studies gave intuitive explanations for various phenomenological signatures; however, they do not provide a structural understanding of the microscopic origin of plectoneme pinning by base-pair mismatches. Finally, the dynamics of plectoneme pinning is yet another aspect that remains unexplored from a theoretical, computational, or experimental standpoint.

In this paper, we develop a computational framework within the OxDNA2 model(47–49) to study plectoneme localization at mismatches. We verify that results obtained in this framework are consistent with all atom simulations of the effects of defects on DNA bending rigidity and reproduce experimental and theoretical features. We use the computational framework to study the effect of force and salt concentration on the localization of plectonemes at mismatches. We find that plectoneme localization is highly reproducible in the high force high salt regime, whereas plectoneme pinning becomes stochastic when the force and salt are lowered. This framework can also be used to study the effect of defects on the dynamics of the supercoiling process and the effect of mismatches on negatively supercoiled DNA.

The use of computational modelling provides molecular-level details of the plectoneme formation process, which are not accessible with existing experimental or theoretical approaches. We establish a computational approach to model the effects of mismatches on DNA supercoiling that offers a complementary approach to statistical mechanical and experimental approaches: the combination of these approaches will permit a comprehensive understanding of DNA mechanics in the presence of defects.

## MATERIALS AND METHODS

We perform molecular dynamics (MD) simulations of a DNA molecule using the OxDNA2 model(47, 48). All the simulations presented in the current paper are performed using the sequence dependent variation of the OxDNA2 model(48) implemented in the LAMMPS(50) simulation software by Henrich et al(49). The OxDNA model has previously been shown to reproduce the behavior of DNA under tension and torsional stress(28) and has also been used to study various structural features of DNA(29, 51–55). In the following sections the simulation parameters are given in the reduced Lennard-Jones (LJ) units, we will provide equivalent parameters in SI units where necessary. The conversion of LJ units to SI units is not simple, specifically because coarse graining eliminates many degrees of freedom, the potential energy surface is flattened, and hence the dynamics are accelerated. Here, the LJ units are converted to SI units using the parameters provided on the OxDNA2 model website and we provide them in the table below.

|  | LJ units | SI units |
| --- | --- | --- |
| Length | $\sigma_{LJ}$ | 0.8518 nm |
| Time | $\tau_{LJ}$ | 3.03 ps |
| Energy | $\epsilon_{LJ}$ | 41.42 pN nm |
| Force | $F_{LJ}$ | 48.63 pN |

### Modelling of mismatched base-pairs

The OxDNA2 model treats each nucleotide as a rigid body with only three interaction sites (base, phosphate, and sugar groups)(47, 48)and as such eliminates many degrees of freedom of individual base pairs compared to all atom simulations, but this coarse-graining approximation is necessary to reach length scales and time scales relevant to the study of super-coiling dynamics of DNA. The OxDNA2 model uses sequence dependent stacking and hydrogen bonding interactions; other interactions between nucleotides are independent of nucleotide type(48). A base pair mismatch in OxDNA2 is simulated by removing the hydrogen bond between the bases. Harrison et al.(56)have previously used the sequence independent version of OxDNA (OxDNA1) to simulate mismatches in a similar manner. Previous studies have shown that many types of mismatched base-pairs have hydrogen bonds(57); therefore, approximating a mismatched base pair without hydrogen bonds is perhaps a crude approximation. However, as we demonstrate below, such an approximation quantitatively reproduces the characteristic decrease in bending stiffness at the mismatch obtained from all atom simulations. Eliminating degrees of freedom through the coarse-graining approach may alter the dynamics of individual base-pairs. But here we are interested in studying the effect of mismatches on the bending rigidity of the local structure of duplex and bending rigidity is not altered due to this approximation (as shown by the favorable comparison with the all atom simulation results, see Fig. S1 in the Supporting Information). Ditmore et al.(18) and Brahmachari et al.(36) have shown that the reduction in local bending rigidity due to the presence of mismatched base-pairs promotes plectoneme pinning.

To calculate the effect of mismatches on the free energy of bending, we simulate a 30 bp long DNA with the mismatch located at the center. We employ umbrella sampling to efficiently sample the free energy landscape. The reaction coordinate (*θ*) is the angle defined between the center of mass of 3 blocks of 5 base pairs (block 1: 4-8 bp, block 2: 13-17 bp, block 3: 23-27 bp). The ends of the DNA are unconstrained. We then apply a harmonic constraint, of type

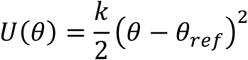

using the COLVARS(58) module in LAMMPS. Here *k* =0.664(*ϵ*_*LJ*_/*rad*^2^). We carry out both forward and backwards sampling of the free energy landscape by first gradually decreasing *θ*_*ref*_ from 180° to 30° in steps of 3°, and then increasing *θ*_*ref*_ from 30° to 180° in steps of 3°. We simulate each step for 3 ∗ 10^4^*τ*_*LJ*_ (90.9 ns) of which the first 750*τ*_*LJ*_ (2.25 ns) are equilibration steps and not considered in subsequent data analysis. The MD simulations are carried out in the NVT ensemble using the Langevin thermostat with damping factors of 1 *τ*_*LJ*_and 10 *τ*_*LJ*_ for the transitional and rotational degrees of freedom, respectively. The equations of motion are integrated using the velocity-verlet algorithm with a timestep of 0.003*τ*_*LJ*_ (9.09 fs). We then extract the free energy associated with bending using the weighted histogram analysis method (WHAM)(59) as implemented by Grossfield (http://membrane.urmc.rochester.edu/?page_id=126).

For the supercoiling simulations, we consider a 610 and 1010 bp long DNA with 50% G-C content and 0, 2, 4, or 6 consecutive mismatches at the center of the DNA. We constrain the 5 base pairs at the top end of the DNA in the x-y plane and also constrain 5 base pairs of the other end in the x-z plane. This allows the DNA ends to freely move in the y and z directions in response to tension and torsional stress while ensuring that the boundary base pairs do not rotate so that the super-helical density remains constant during the simulation. To ensure that the nucleotides don’t pass around the DNA ends, we apply a purely repulsive harmonic potential of type *U*(*Z*) = *k*(*Z* − *Z*_*ref*_)^2^ that acts in the x-y plane and repels all but the boundary base pairs if they move beyond the boundary base pair in the z direction. Here *k* = 1000(*ϵ*_*LJ*_/*rad*^2^) and *Z*_*ref*_ is 5σ_*LJ*_away from the last base pair at each end.

We define the equilibration time *τ*_*equi*_using the autocorrelation function of the DNA end-to-end distance (*R*_*end*_). The auto-correlation function of *R*_*end*_,(29) is calculated as

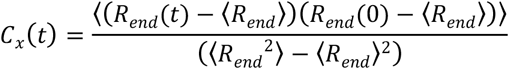

The equilibration time is then given by

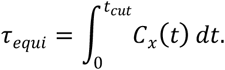

Here the upper limit of the integral *t*_*cut*_ is selected as the time when *C*_*x*_(*t*) = 0. It is necessary to use this limit to eliminate the effect of the noisy tail of *C*_*x*_(*t*). To collect data, we discard the first 2 ∗ *τ*_*equi*_timesteps. In the current work, all the simulations have been carried out for at least 12 ∗ *τ*_*equi*_.

We start each simulation from an equilibrated condition. We apply torsional stress by rotating the top end of the DNA around the z axis for fixed number of turns (*N*_*turns*_). To select the value of *N*_*turns*_, we first conducted preliminary simulations and then selected the smallest value of *N*_*turns*_ for which we could obtain a stable plectoneme within a reasonable simulation time. A larger value of *N*_*turns*_ would produce a larger plectoneme that would stabilize faster and hence require less simulation time. On the other hand, a larger plectoneme will restrict the motion of the plectoneme along the DNA backbone and hence it would require a longer simulation to accurately capture the entropic effects of plectoneme diffusion along the DNA backbone.

To apply tension, we apply a force along the z-axis to the top base-pair. For all simulations we use the Langevin C(60) thermostat with a timestep of 0.01*τ*_*LJ*_, and unless otherwise specified, damping factors of 5 *τ*_*LJ*_ and 10 *τ*_*LJ*_ for the transitional and rotational degrees of freedom respectively.

For simulations at a salt concentration of 1 M, we performed 5 simulations (using different random numbers as the seed for the Langevin thermostat) of a 610 bp long DNA for each mismatch value. Simulations were performed for 2.6 × 10^7^*τ*_*LJ*_ (78.78 *μs*) for the case of 0 mismatches; 2.7 × 10^7^*τ*_*LJ*_ (84.08 *μs*) for 2 mismatches, and 2 × 10^7^*τ*_*LJ*_ (60.6 *μs*) for 4 and 6 mismatches. For 4 and 6 mismatches, we used a damping factor of 15 *τ*_*LJ*_ for the rotational degrees of freedom.

For simulations at a salt concentration of 0.2 M, we performed simulations of 610 bp and 1010 bp DNA molecules. For the 610 bp DNA, we performed 20 distinct simulations for each mismatch size for approximately 3. 5 × 10^7^*τ*_*LJ*_ (105 *μs*) using a timestep of 0.01 *τ*_*LJ*_ and a damping factor of 15 *τ*_*LJ*_ for the rotational degrees of freedom.

For the 1010 bp DNA we performed 10 simulations for each mismatch. The simulations were run with a timestep of 0.01*τ*_*LJ*_ for approximately 3.710^7^*τ*_*LJ*_(112.11*μs*).

Combined these simulations required a total of 5.2 × 10^6^ CPU hours.

### Plectoneme detection algorithm

To detect a plectoneme we used a modified version of the algorithm provided by Matek et al.(28)The plectoneme detection algorithm is as follows:

- Find the center of mass of each base pair (bp) in the DNA.
- Start at the bottom bp and loop over all bp.
- Find the distance between the current bp (*i*) and all other base pairs (*j*) beyond a cut-off of *N*_*c*_ bp along the DNA backbone, i.e., distance between bp *i* and bp *j*, for all *j* > *N*_*c*_.
- If any distance is less than *d*_*cuttoff*_; the bp is identified as the beginning of plectoneme.
- To detect the end of plectoneme we utilize the concept of contact points. Once the beginning of the plectoneme is detected, we skip the next *Nc* bp’s along the backbone and identify the bp closest to the bp identified as the start of the plectoneme. This bp is defined as the plectoneme end.
- The plectoneme center is defined as the mean of the bp indices of the beginning and the end of the plectoneme.

We typically choose *N*_*c*_ = 80, and 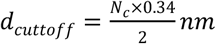. The algorithm allows us to detect multiple plectonemes, but the size of each plectoneme detected is larger than *N*_*c*_. We also verified that changing *N*_*c*_ does not significantly alter the results.

### Calculation of the end loop angle

The end loop angle is defined between the center of mass of 3 blocks of 5 bp. If the center of the plectoneme is considered as base *i*, the 3 blocks would be block 1: *i*−12 *to i*−8 bp, block 2: *i*−2 *to i*+2 bp, block 3: *i*+8 *to i*+12 bp.

We also note that our plectoneme detection algorithm does not always provide the exact center of the plectoneme. For the case of high force and high salt concentration, where the plectoneme center is pinned at the location of the mismatch, the plectoneme detection algorithm calculates the center of the plectoneme with an error of around ± 10 bp. Keeping this in mind, to calculate the angle of the end loop we vary the center of the plectoneme between *i* ± 15 bp and select the lowest angle as the angle of the end loop.

### Calculation of twist

To calculate the twist between two consecutive base pairs, we use the method defined by Skoruppa et al. (61) and employed in subsequent studies (55, 62); specifically the definition of *Triad II*, as defined by Skoruppa et al., is used for all calculations.

Briefly, the twist for each bp step can be calculated by defining two vectors, one vector connecting the centers of mass of two nucleotide in a bp 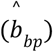 and the second vector connecting two base pairs along the DNA contour 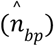. Twist can then be defined as the angle between the two 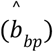 for each base pair in the base pair step along the 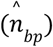. A more detailed description can be found in Skoruppa et al(61).

### Calculation of writhe

To calculate the writhe, we carried out simulations of DNA under tension but no torsional stress. We analyzed these simulations to obtain *Lk*_0_ = *Tw*_0_, which is the twist inherent in the DNA with different number of mismatches.

Writhe in DNA is then defined as:

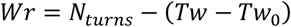

where, *Tw* is the average twist in DNA under tension and torsional stress.

### Calculation of diffusivity

To obtain the diffusivity of the plectoneme we first calculate the mean squared displacement (MSD) of the plectoneme center *P*_*cent*_.

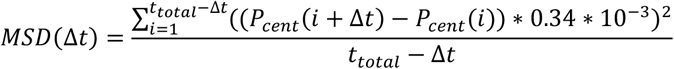

We calculate MSD for Δ*t* ∈ {0,20}frames, where Δ*t* is the time lag, we select trajectories that have at least 400 continuous plectoneme centers. We then average all the MSD curves for a given condition (see Fig. S3 in the Supporting Information).

The diffusion constant can be obtained from the MSD curve using the relation *MSD* = 2*Dt*, we perform linear fitting for the first 8 points on the MSD vs time curve to obtain the diffusion constant.

### Visualization

Visualization of the simulation configurations is carried out using the OVITO(63) simulation software. LAMMPS provides the center of mass and orientation of each nucleotide in the form of a quaternion. We post-process this information using MATLAB to find the location of base interaction sites and the backbone site for all nucleotides. The process used is similar to the process employed in the visualization code provided by Oliver Henrich with the OxDNA module in LAMMPS.

## RESULTS AND DISCUSSION

### Coarse-grained estimates of the effects of DNA defects on bending free energy

Coarse-grained simulations of DNA, in particular OxDNA2, afford the requisite trade-offs between molecular-level details and computational speed and efficiency to permit millisecond-scale simulations of kilobase length DNA(29). OxDNA has been successfully applied to study DNA supercoiling and has been benchmarked against single-molecule measurements of DNA supercoiling(28). For these reasons we decided to develop a mismatch model of DNA in the OxDNA2 framework. We first validated the effects of defined mismatches on DNA bending elasticity for OxDNA2. Fig. 1(a,b) shows the variation of the total Gibbs free energy (Δ*G*) of the DNA molecule as a function of the bend angle imposed at the centre of the DNA molecule containing different numbers of consecutive mismatches at the centre of the DNA for high (Fig. 1 a) and low (Fig. 1 b) salt concentrations. A bending angle of *θ*_*b*_ =170° corresponding to a nearly straight DNA, is the lowest free energy configuration of DNA with and without mismatches. Previous studies(56, 64, 65) have also found the lowest free energy configuration to be at an angle slightly less than 180° (i.e. a completely straight segment of DNA). Bending at the DNA centre reduces this angle. The presence of mismatches reduces the bending energy of the DNA at the location of the mismatch(39). Accordingly, Δ*G* increases from the minimum at *θ*_*b*_=170° as the included angle decreases for all DNA molecules, but for a given bend angle the energy progressively decreases with increasing mismatch size, from a maximum for no mismatch. This data, establishing the effect of mismatch size on bending energy, is consistent with results from previous studies(18, 36, 39) and suggests that the mismatch model is feasible for investigating DNA plectoneme formation in the presence of mismatches through coarse-grained approaches.

**Figure 1.**
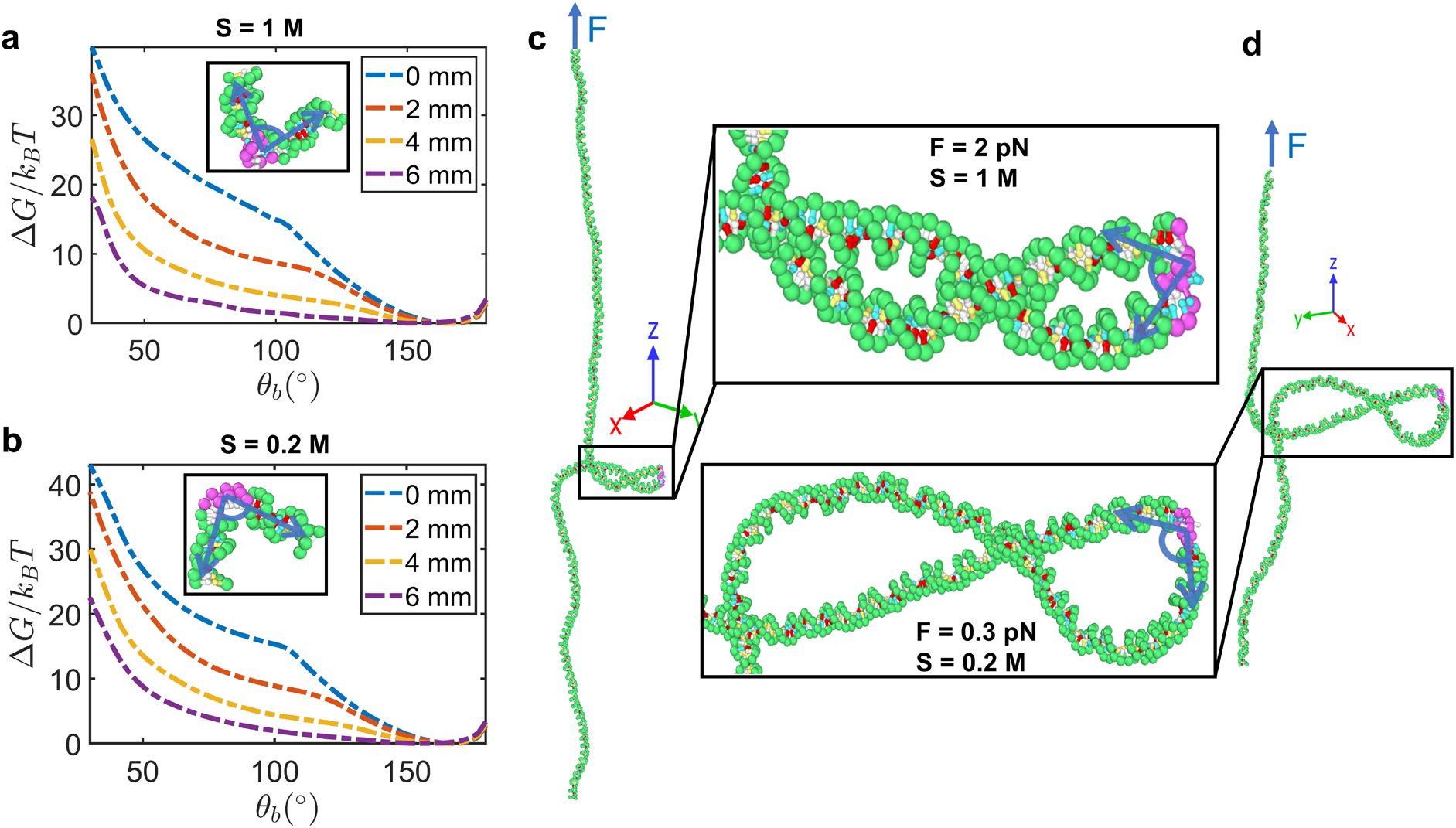
DNA mismatches lower bending energy and nucleate plectonemes in supercoiled DNA. **(a, b)** Free energy of a 30-bp DNA molecule as a function of the bending angle *θ*_*b*_ (in degrees) imposed at the center of the DNA (i.e., the 15^th^ bp) (see also the methods section). The results are shown for (a) high salt concentration (1 M monovalent salt) and (b) low salt concentration (0.2 M monovalent salt). For both cases, there is no external stretching force applied to the DNA and the results are shown for different numbers of consecutive G:T mismatches, introduced at the center of the DNA. The insets illustrate the DNA configuration (with 6 consecutive G:T mismatches introduced at the center of the DNA) corresponding to a bending angle of approximately 95°, indicated by the blue arrows. Here the green spheres represent the sugar group of the intact base pairs whereas pink spheres represent the sugar group of the mismatched bases. Red, yellow, white, and blue spheres represent the A, C, G and T base groups, respectively. **(c, d)** Simulation snapshots of DNA plectonemes formed by positively supercoiling (over-winding) a 610-bp DNA molecule with 6 mismatched bases under the conditions of (c) High salt (1 M monovalent salt) and high force (F = 2 pN) and (d) Low salt (0.2 M monovalent salt) and low force (F = 0.3 pN). The DNA is subjected to a constant force one end, while the other end is fixed. The torque generated in twisted DNA (3.46 and 2.31 turns in (c) and (d), respectively) results in buckling of the DNA to form a plectoneme. Expanded views of the plectonemes along with the definition of the bend-angle at the mismatch location (blue arrows) are shown in the insets of (c) and (d). A detailed description of the end loop bend angle can be found in the methods section. For the sake of brevity, mismatches are referred to as mm in the legends.

We next performed simulations of positively supercoiled DNA (610 bp and 1010 bp) containing 0, 2, 4, or 6 G:T mismatches at the center of the DNA molecule (Fig. 1 c and 1 d). To compare the simulation results with experimental and theoretical results, simulations were run with two different conditions of monovalent salt and force: high monovalent salt (1 M) and high force (2 pN); and low monovalent salt (0.2 M) and low force (0.3 pN) (Fig. 1 c and 1 d). We refer to the conditions of 1 M monovalent salt and 2 pN of force as the high force - high salt condition, and conditions of 0.2 M monovalent salt and 0.3 pN as the low force - low salt condition. We have summarized the different simulation condition in Table 1. The most important feature distinguishing the plectonemes formed in these two cases is the angle of the end-loop (defined in the insets of Fig. 1 (c, d) and in the methods). The end-loop angle is smaller under the high force and salt condition, indicating a larger extent of deformation or kinking. For example, in the snapshots shown in Fig. 1 (c, d), this angle is 46.8° and 143.3° for the high and low force and salt conditions, respectively.

**Table 1.** Details of simulation conditions.

| DNA Length (bp) | Force Applied (pN) | Implicit Monovalent Salt concentration (M) | Turns applied ($\Delta Lk$) | Referred to as |
| --- | --- | --- | --- | --- |
| 610 | 2 | 1 | 3.46 | High force - High salt |
| 610 | 0.3 | 0.2 | 2.31 | Low force - Low salt |
| 1010 | 0.3 | 0.2 | 2.87 | Low force - Low salt |

Kinking the end-loop brings similarly charged segments of the DNA closer to each other and contributes an additional electrostatic free energy cost to plectoneme formation. Under high salt conditions, the electrostatic repulsion is screened over a much smaller distance leading to a reduced Debye length. This, in turn, reduces the electrostatic repulsion between the buckled segments and hence reduces the energetic cost of buckling. Furthermore, the external force imposes a second energetic cost associated with buckling related to the work done against the force in decreasing the extension of the DNA by an amount equal to the loop size. As a result, smaller loops with increased kinking are favored under higher applied forces. Previous theoretical and experimental studies obtained similar force and salt concentration dependent increases in the kinking of the end loop(66). Our results demonstrate that kinking is enhanced in the presence of mismatches, but that the extent of kinking is governed by the applied force and salt concentration.

### Mismatches enhance plectoneme pinning

A key question related to the supercoiling of DNA containing mismatches is the degree to which the mismatch localizes or pins the plectoneme by stabilizing a sharp bend at the plectoneme tip. This pinning effect involves both the degree of bending or kinking at the plectoneme tip and the probability of the mismatch being located at the tip of the plectoneme. Consistent with previous studies(66), we observe a decrease in the end loop angle between the high force - high salt and low force - low salt conditions. For both the simulation conditions, we observe a decrease in the end loop angle (when the plectoneme center is pinned at the mismatches) with increasing number of mismatches (Fig. 2 d-f). This is consistent with the decrease in the local bending energy in proportion to size of the mismatch (Fig. 1 a and b). This decrease in local bending energy results in a decrease in the end loop angle (increased kinking) when the mismatch is located at the tip of the plectoneme (operationally defined by the centre of the plectoneme coinciding with the mismatch, see below). Previous experimental studies (67) have shown that the intrinsic bend of a DNA segment can affect plectoneme pinning. We characterized the intrinsic bend of the DNA sequence used here (see section S6 and Fig. S5 in the Supporting Information) to ensure that the specific DNA sequence does not affect the plectoneme pinning probability. We note that the ability of the OxDNA2 model to reproduce intrinsic bends in DNA sequences is not well characterized. Nonetheless, the data in supplementary Fig. S5 provide a qualitative estimate of sequence tracts with strong intrinsic bends.

**Figure 2.**
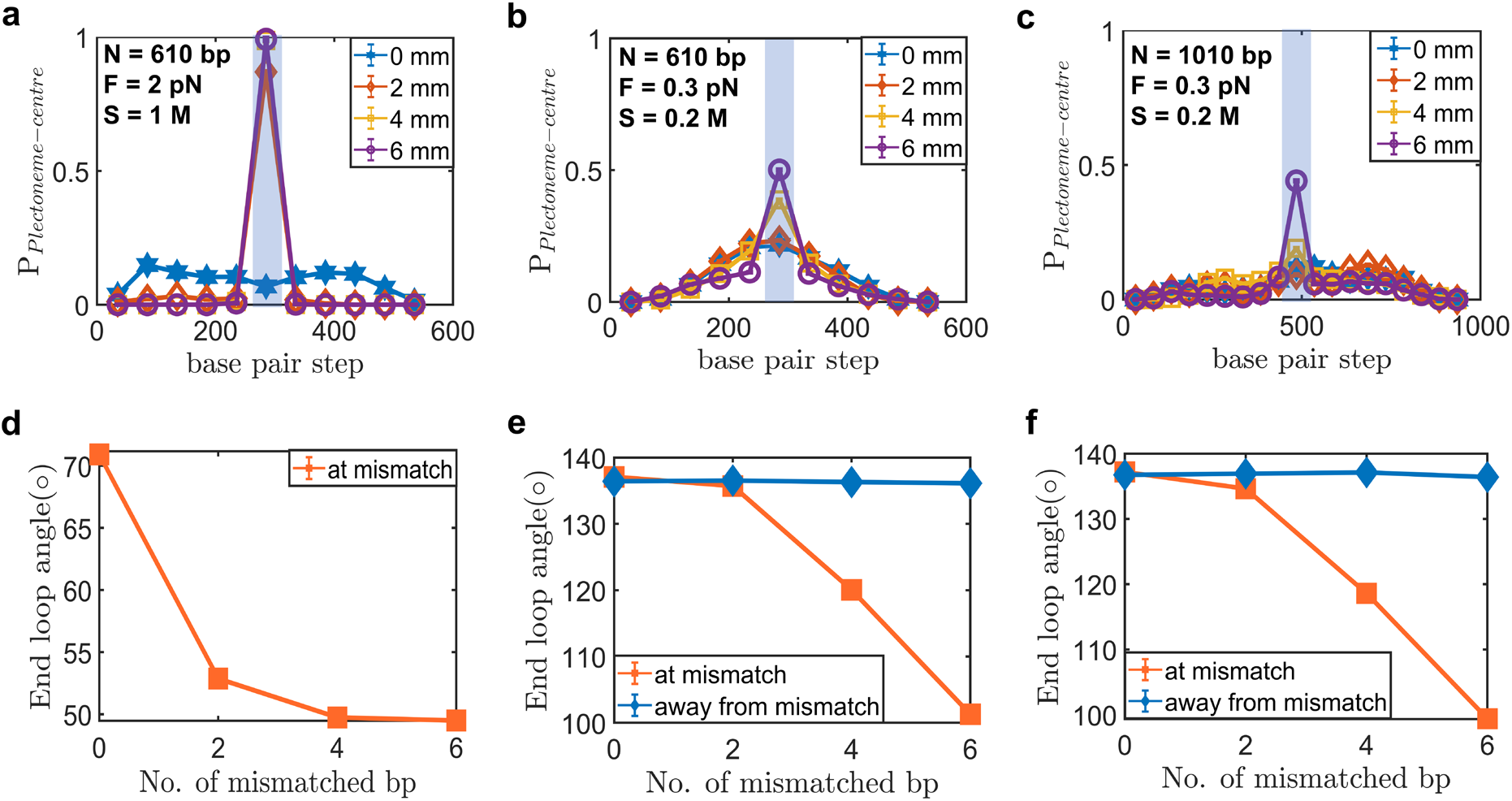
Plectoneme end loop kink angle and position. **(a-c)** Probability distribution of the plectoneme center position. Probability of finding the plectoneme center (P_Plectoneme-center_) at a given DNA base pair for (a) 610 bp DNA with F=2 pN and 1 M monovalent salt, (b) 610 bp DNA with F=0.3 pN and 0.2 M monovalent salt, and (c) 1010 bp DNA with F=0.3 pN and 0.2 M monovalent salt. The location of the DNA mismatch is denoted by the blue shaded region. Error bars (standard error of the mean, SEM) are smaller than the symbol size. **(d-f)** Average end-loop angle in degrees (see the details of the averaging procedure in the Methods section) of the DNA plectoneme [this angle is defined by blue arrows in the insets of Figs. 1 c and Figs. 1 d] as a function of mismatch size (in base pairs) for (d) 610 bp DNA with F=2 pN and 1 M monovalent salt, (e) 610 bp DNA with F=0.3 pN and 0.2 M monovalent salt, and (f) 1010 bp DNA with F=0.3 pN and 0.2M monovalent salt. The orange line represents the average end loop angle when the plectoneme center is pinned at the mismatch, whereas the blue line represents the average end loop angle when the plectoneme center is not pinned at the mismatch. Error bars (SEM) are smaller than the symbol size. For the sake of brevity, mismatches are referred to as mm in the legends.

The probability of the mismatch being localized at the tip of the plectoneme depends on the DNA length, applied force, monovalent salt concentration, and mismatch size. Under the high force - high salt condition (Fig. 2 a), the center of the plectoneme always coincides with the location of the mismatch for 2, 4, and 6 mismatches. This observation is in agreement with the single molecule magnetic tweezers experiments conducted by Dittmore et al.(18) and results of previous simulation study(68). Furthermore, the DNA is increasingly sharply bent at the mismatch as the number of mismatched bases increases [Fig. 2 d]. However, for the low salt - low force condition there is a non-unity (<1) probability that the mismatch is located at the centre of plectoneme, though this probability rapidly increases with the number of mismatches [Fig. 2 b and Fig. 2 c]. This result has been predicted theoretically(36). The finite probability of buckling at the mismatch for the low salt and low force conditions leads to a bifurcation of the end loop angle: the end loop angle is constant, independent of the mismatch size, when the mismatch is not located at the tip of the plectoneme but decreases significantly with increasing mismatch size when the mismatch coincides with the tip of the plectoneme (Fig. 2 e and 2 f). The difference in the plectoneme localization between the two conditions (high-salt-high-force and low-salt-low force) can be explained through a free energy argument. Brahmachari *et al.* (36) developed quantitative expressions relating the probability of plectoneme buckling and pinning to the relative entropic and enthalpic contributions to the free energy in the presence and absence of mismatches. Here we provide qualitative energy arguments to provide physical insight into the observed buckling and pinning behavior.

For the high force - high salt case, bending or kinking at the mismatch is favored [Fig. 2 a]. Fig. 1 (a) indicates that sharper bending requires higher energy. Higher forces lead to increased bending of the end loop since this minimizes the extent of the plectoneme loop, which in turn minimizes the change in extension of the DNA and the corresponding work against the external force. In this scenario, it is energetically favorable if the bending occurs at the location of the mismatch where the bending rigidity is decreased. However, there is an entropic cost of buckling at the mismatch (equivalent to localizing the plectoneme at the mismatch), which limits the possible configurations of the plectoneme as compared to the case where the plectoneme can form at any location on the DNA molecule. Mathematically, we can write this as:

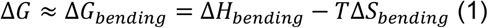

where Δ*G*, Δ*H*, and Δ*S* are the changes in the Gibbs free energy, the enthalpy, and the entropy, respectively.

For the case of high salt - high force, the plectoneme is pinned at the location of the mismatches leading to a large loss of entropy. Therefore, we can split the energy contributions as follow:

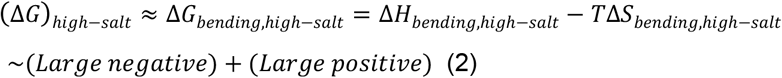

The equilibrium is driven by the competition between a large favorable bending enthalpy (due to the localization of the bending deformation at the defect) and a large unfavorable bending entropy (due to the pinning of the plectoneme). In the current work, we find that for a force of 2 pN applied to a 610 bp DNA with 2, 4 and 6 consecutive mismatched base pairs, the enthalpic gain due to bending at the mismatch overcomes the entropic loss due to pinning; hence the plectoneme is always pinned at the mismatch.

Next, we consider the free energy for the low salt - low force condition. Here too, we can write the free energy change as:

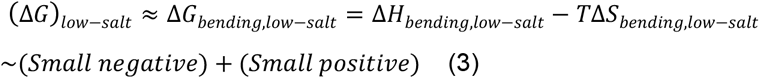

Here both the enthalpy and the entropic components are smaller in magnitude. The equilibrium is driven by the competition between a weakly favorable bending enthalpy (due to a higher bending angle of the end loop, the enthalpic difference for bending at the defect for different numbers of mismatches is lower as compared to the high force case, see Fig. 2 e and Fig. 2 f) and a weakly unfavorable bending entropy (due to a larger plectoneme for the low force-low salt case compared to the high force-high salt case, the entropic cost of pinning is lower for a given length of DNA, see Fig. 4). In the case of low force - low salt for a 610 bp DNA, we find an increase in probability of plectoneme pinning with increasing number of mismatches. To probe the effect of the entropic contribution to the pinning probability, we studied a 1010 bp DNA with 0, 2, 4 and 6 bp mismatches under the same conditions. Pinning in a longer DNA results in a larger entropic loss due to the presence of a larger number of configurations arising from plectoneme diffusion. Consistent with this reasoning, we find that for the 1010 bp DNA, the enthalpic gain due to bending at the mismatch is able to overcome the entropic loss and lead to plectoneme pinning only for the case of 6 mismatches (Fig. 2 f).

**Figure 4.**
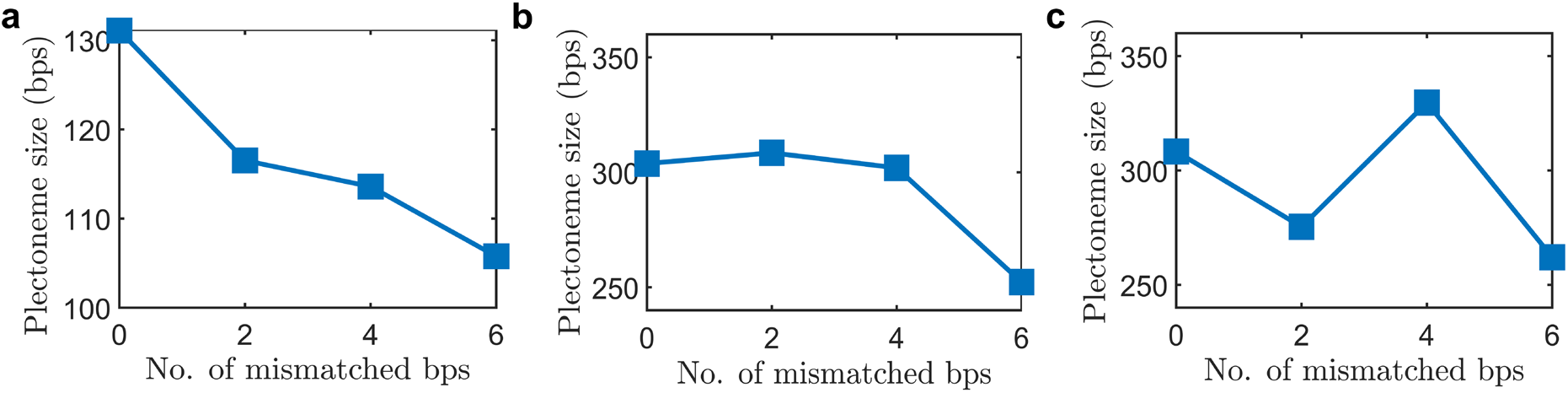
Influence of mismatches on the length of DNA in plectoneme. Average size (in bp) of the length of DNA in the plectoneme (see Methods section for details of the averaging procedure) as a function of the number of consecutive mismatched base pairs introduced at the DNA center for (a) 610 bp DNA with F=2 pN and 1 M monovalent salt, (b) 610 bp DNA with F=0.3 pN and 0.2 M monovalent salt, and (c) 1010 bp DNA with F=0.3 pN and 0.2 M monovalent salt. Note the differences in the y-axis scales. Error bars (SEM) are smaller than the symbol size.

### Simulation results of plectoneme pinning agree with statistical mechanical model

To verify the results of the MD simulations, we compared them with results obtained from a previously published theoretical model (36) that is built upon the statistical mechanical behavior of dsDNA as a semiflexible polymer. The model explicitly incorporates various free-energy components, such as the contribution from DNA bending associated with the end loop and the plectoneme, the Debye-Hückel electrostatic energy associated with the plectoneme wrapping, and the stretching energy under an external force. The presence of a defect or base-pair mismatch is theoretically modeled as a mismatch-size dependent reduction in the energy component corresponding to DNA bending in the end loop. This energy reduction is incorporated through a defect size parameter, ε, which varies from 0, for no mismatches, to a maximum of 1 as the extent of the mismatch increases. The energy and size of the plectoneme end loop are rescaled by (1− ε). With this simplifying assumption, the energetic differences associated with a mismatch are lumped together in a single factor, which is likely an oversimplification of the underlying energetic considerations associated with mismatches. Brahmachari et al.(36) were able to provide rough estimates for the relation between the number of mismatches and the defect size parameter or the decrease in bending energy through comparisons with experimental measurements of defect facilitated DNA buckling(18).

Current simulations allow detailed microscopic understanding of the bending energy reduction associated with mismatches. Comparing the structure of the end loop and the energy associated with a kinked end loop, we obtain values corresponding to the defect-size parameter for different mismatch sizes (see table S1 in the Supporting Information). Interestingly, the defect size parameter depends on the force and salt conditions. This suggests that the thermodynamic size of the defect, which is the parameter that controls the probability of buckling at the defect site, is not only dependent on the size of the mismatch but also depends on the force and the salt conditions. Using these values for the defect size, we find that the theoretical model predictions are in reasonable agreement with the simulation results, showing the same trends (see Fig. S4 in the Supporting Information). More specifically, for the high force and high salt conditions the simulations and the theoretical calculations quantitatively agree, whereas for the low force and low salt conditions (probabilistic regime) the theoretical probability of plectoneme pinning is over-estimated compared to the simulation results. The free energy model treats thermal fluctuations as a perturbation about the highly extended DNA state, which could possibly underestimate the effects of fluctuations in the low-force regime.

We plan to conduct experiments in the future to accurately probe the probabilistic regime of buckling at the mismatch that will allow us to provide experimental constraints for the modeling and simulation approaches.

### Mismatches absorb twist but do not affect writhe

In addition to providing the location and bending angle of the tip of the plectoneme, the MD simulation approaches provide detailed information concerning every base pair in the DNA molecule. An important consideration that has not been addressed in prior experimental or theoretical studies is the effect of the mismatches on the torsional compliance of the DNA molecule. To address this, we calculated the average twist over each base-pair for both the intact and mismatch bases in each simulation (Fig. 3 a-c). The presence of mismatched bases is expected to cause a decrease in the local torsional stiffness. This effect can be seen clearly for the high force - high salt case [Fig. 3 a]: the twist per bp step at the mismatch is higher than the twist per bp step at the intact DNA, and it increases with the size of the mismatch region, whereas the twist per bp step for the intact DNA remains constant. For the low force-low salt case, there does not appear to be a similar reduction in torsional stiffness [Fig. 3 b and 3 c)]. Curiously, compared to the intact DNA, the twist per bp step at the mismatch actually decreases for the case of 2 mismatches for both 610 bp [Fig. 3 b] and 1010 bp [Fig. 3 c], but the twist per bp step increases for the 4 and 6-mismatch cases. We note that the twist at the 2 bp mismatch will be affected more by the flanking base pairs as compared to the 4 and 6-mismatch cases, which might be responsible for a lower twist at the mismatch.

**Figure 3.**
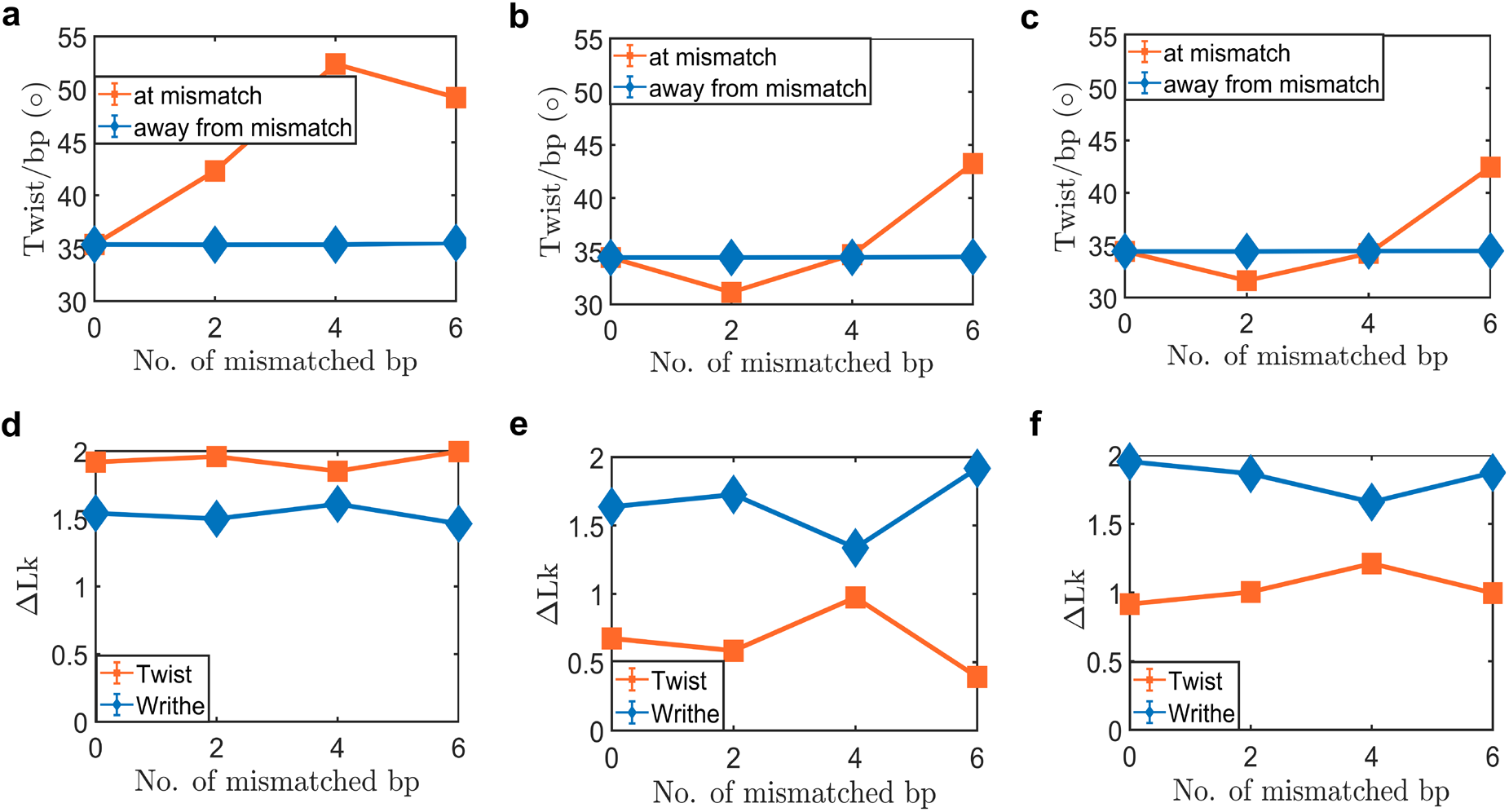
Twist and writhe in supercoiled DNA containing mismatches. **(a-c)** Average twist per bp step in degrees (The details of the averaging procedure are provided in the Methods section) of the DNA as a function of the number of consecutive mismatched base pairs introduced at the DNA center for (a) 610 bp DNA with F=2 pN and 1 M monovalent salt, (b) 610 bp DNA with F=0.3 pN and 0.2 M monovalent salt, and (c) 1010 bp DNA with F=0.3 pN and 0.2 M monovalent salt. Orange line represents the average twist per bp step at the mismatch, Blue line represents the twist per bp step averaged over all the intact base pairs. Error bars (SEM) are smaller than the symbol size. **(d-f)** Distribution of twist and writhe as a function of the number of mismatches for (d) 610 bp DNA with F=2 pN and 1 M monovalent salt, (e) 610 bp DNA with F=0.3 pN and 0.2 M monovalent salt, and (f) 1010 bp DNA with F=0.3 pN and 0.2 M monovalent salt. The details of the averaging procedure can be found in the Methods section. Error bars (SEM) are smaller than the symbol size.

To better understand the twist accumulation at the mismatch, we performed simulations of torsionally free and stretched DNA with the same sequence as the DNA under torsional stress (see Fig. S2 in the Supporting Information). In torsionally free DNA, we observed the same decrease in the twist at the mismatch step for the 2-mismatch case and subsequent increase in twist for 4 and 6 mismatches. This would suggest that for the 2-mismatch case, the twist at the mismatch is lower even when no torsional stress is applied to the DNA (see Fig. S2 in the Supporting Information). When torsional stress is applied, we observe a higher twist accumulation at the mismatch as compared to that at the intact base pairs due to torsional softening. The effect of torsional softening can be clearly seen for the high salt high force case [Fig. 3 a]: here the twist at the mismatch is larger than that at the intact bases in the duplex. The twist at the mismatch for the low salt low force case is similar to the case with no torsional stress, likely due to the lower applied torsional stress. All atom molecular dynamics simulations have been performed to quantify the effect of a mismatch on the twist per bp step distribution in DNA (57, 69). These studies found that for the G:T mismatch used in the current study, the twist at the mismatch is similar to the twist at the intact DNA. To the best of our knowledge, there has been no study relating the number of consecutive mismatches to the twist per bp step distribution in the DNA.

We next probed possible global effects of the mismatches on the distribution of twist and writhe in the supercoiled DNA molecules. For all conditions and mismatch sizes, we find the twist and writhe stored in the DNA are not significantly altered by the presence of mismatches (Fig. 3 d-f). In the current work, the mismatches comprise a small fraction of the DNA (a maximum of 6 mismatches in a 610 bp DNA), hence the torsional softening at the mismatches does not significantly affect the twist, and hence the writhe, stored in the DNA. However, for the low force - low salt conditions (Fig. 3 b and 3 c), we see a slight increase in the twist stored in the DNA for the 4-mismatch case. This increase is consistent for both the 610 bp and 1010 bp case and may be statistically significant. A future study of how the twist for the 4-mismatch case varies with increasing torsional stress will be helpful in determining the significance of this observation.

### Mismatches allow DNA to accommodate the same writhe with a smaller plectoneme

Whereas the degree of bending or kinking at the tip of the plectoneme is an important determinant of the localization of the plectoneme that depends critically on the size of the mismatch in addition to the salt and force conditions, it is not readily experimentally measurable. Conversely, the extent of the DNA molecule that is in the plectoneme can be accurately measured experimentally since it corresponds to the decrease in DNA extension associated with plectoneme formation. Fig. 4 provides the average size of the DNA plectoneme (length of DNA in the plectoneme) as a function of the number of consecutive bp mismatches introduced at the DNA center for different combinations of the applied force (F), salt concentration (S), and DNA length (N). The plectonemes are smaller under the high force - high salt condition as compared to the low force - low salt condition. This is also evident from the snapshots provided in Fig. 1. The writhe in the DNA is not affected by the presence of mismatches (see Fig. 3 d-f), yet we see a decrease in the plectoneme size with increasing number of mismatches. The presence of the mismatch causes a decrease in the end loop angle [Fig. 2 d-f] and a tighter plectoneme end loop reduces the plectoneme size while not affecting the writhe. Although we note that the decrease in plectoneme size is not universal and seems to disappear for the longer plectoneme.

The relation between bend angle (Fig. 2 d-f) and plectoneme extent (Fig. 4) provides an indirect approach to estimate the bend angle from the experimental quantification of the changes in DNA extension as the plectoneme forms.

### Mismatches reduce plectoneme diffusivity

While the statistics of plectoneme formation and conformation permit comparison with experiments, the simulation results provide a wealth of additional information concerning plectoneme dynamics (Fig. 5 a-d). Tracking the position of the plectoneme along the DNA molecule over time provides a dynamic view of the motion and the pinning of the plectoneme as a function of the size of the mismatch region under different force and salt concentration conditions.

**Figure 5.**
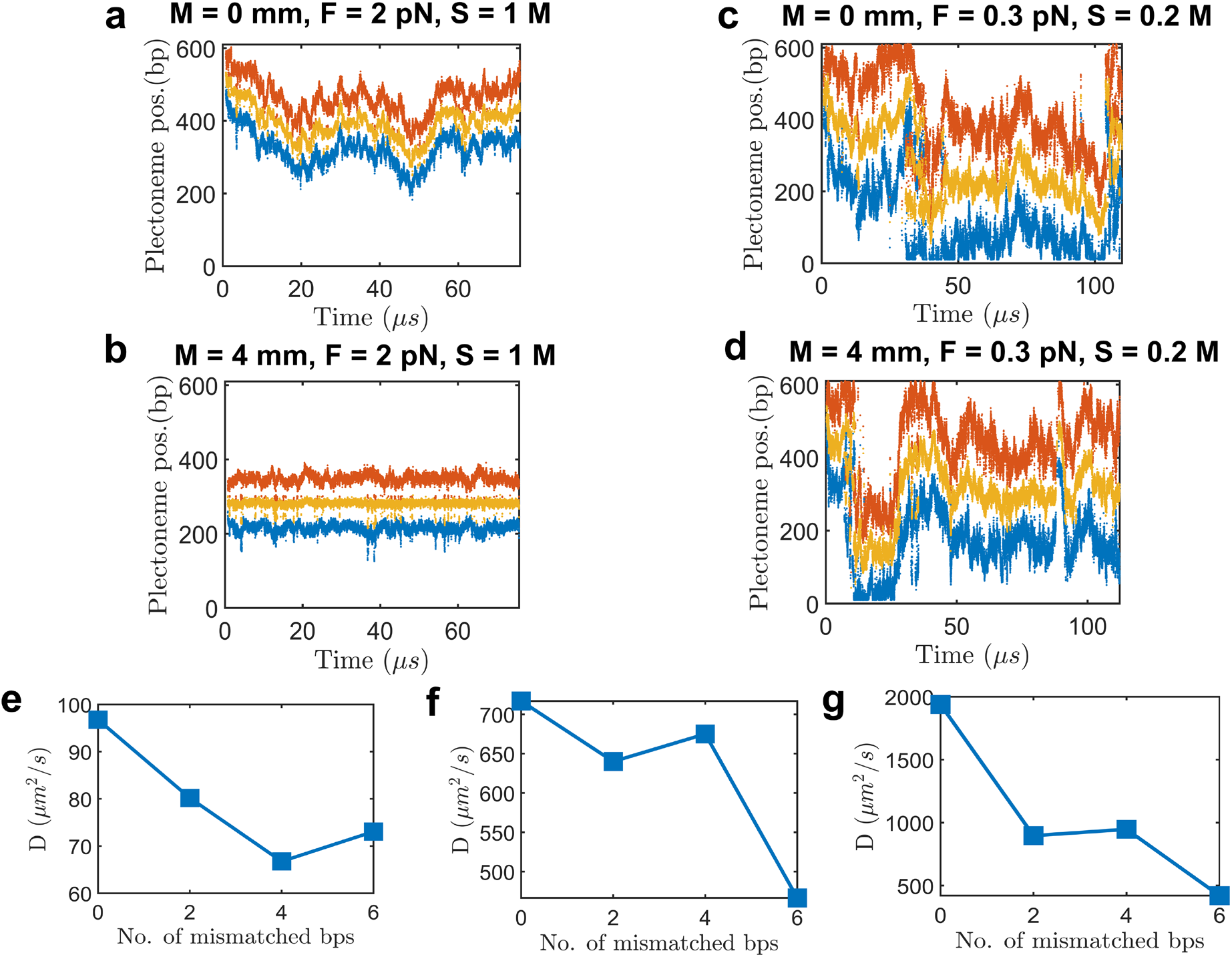
Mismatches slow plectoneme dynamics. **(a-d)** Time-dependent location of the beginning (blue line), center (yellow line), and end (red line) of the plectoneme for (a) high salt and high force with intact DNA (M=0), (b) high salt and high force with 4 consecutive mismatches (M=4) at the DNA center, (c) low salt and low force with intact DNA (M=0), and (d) low salt and low force with 4 consecutive mismatches (M=4) at the DNA center. **(e-g)** Diffusivity of plectonemes as a function of the number of mismatches for (e) 610 bp DNA with F=2 pN and 1 M monovalent salt, (f) 610 bp DNA with F=0.3 pN and 0.2 M monovalent salt, and (g) 1010 bp DNA with F=0.3 pN and 0.2M monovalent salt. For the sake of brevity, mismatches are referred to as mm in the legends.

For intact DNA, the plectoneme is significantly less mobile under the high force - high salt condition than the low force – low salt condition (Fig. 5 a and 5 c). For DNA with four mismatches the plectoneme is strongly pinned at the mismatch location under the high force - high salt conditions [Fig. 5 b]. This is reflected in the unity probability of the plectoneme being centered at the mismatch location [Fig. 2 a]. Conversely, for DNA containing four mismatches under low force - low salt conditions, the plectoneme remains highly mobile, indicating a lack of pinning of the plectoneme at the mismatch. This is reflected in the low, significantly less than unity, probability of the plectoneme being centered at the mismatch location under low salt and low force conditions [Fig. 2 b and 2 c].

To compare the mobility of the plectoneme under different conditions, we calculated the diffusion constant of the plectoneme center (Fig. 5 e-g). The mean squared displacement curves used to obtain the diffusion constant are provided in the supplementary information (see Fig. S3 in the Supporting Information). For plectonemes in both the high force-high salt and the low force-low salt regime, there is a decrease in the plectoneme mobility with increasing number of mismatches (Fig. 5 e-g). The diffusion constant for intact DNA is similar to the value obtained by Matek et al.(28) However, the diffusivity obtained in the current work is 3 orders of magnitude larger than the experimentally observed diffusion constant. In the experiments performed by van Loenhout et al.(70), the diffusion constant is calculated for plectonemes larger than 4 kb. In the current study, the average plectoneme size is considerably smaller (Fig. 4), a smaller plectoneme will be more mobile. Another factor that may increase the apparent diffusivity is the use of a coarse-grained model with implicit solvent in the current work. Coarse-graining decreases the degrees of freedom in the system and hence accelerates the dynamics. Similarly, the use of implicit solvent decreases the effective viscosity, which further accelerates the conformational dynamics. As a result, the plectoneme diffusivity data provided here represents the qualitative trend of the effect of mismatches on plectoneme diffusivity, but the diffusion constants are likely over-estimated. It is possible to decrease the diffusivity of the plectoneme, but this would result in inefficient sampling. The simulation time is limited to the order of micro-seconds, and a higher diffusivity coefficient allows the simulation to sample phenomenon observed in experiments.

## CONCLUSIONS

To the best of our knowledge these results represent the first simulation-based evidence of the manner in which the interplay of force, salt concentration, and DNA size affects the localization of a supercoiled plectoneme in a DNA containing mismatches. We find that in the physiological regime of monovalent salt concentration and force, entropy plays an important role in preventing plectoneme pinning. As a result, under physiological conditions, small mismatch defects position plectonemic domains probabilistically; larger mismatches lead to gradually more deterministic positioning. Although these results were obtained for positive supercoiling, so they could be compared with existing experimental and theoretical results, they are nonetheless applicable to important physiological processes including transcription and replication that generate positive supercoiling *in vivo*. Furthermore, DNA damage repair is known to be coupled with both replication and transcription and the results we provide could potentially indicate a physical mechanism coupling these processes to DNA damage recognition via positive supercoiling. More generally, we provide a coarse-grained simulation approach to investigate the effects of mismatches on DNA supercoiling and plectoneme formation that can provide details of the conformations, structures, and dynamics of the system that are not accessible from analytical theories or current experimental approaches.

In the future we plan to study negatively supercoiled DNA with mismatches using the framework described here. We also plan to experimentally validate the results obtained here.

## SUPPLEMENTARY DATA

Supplementary Data are available at NAR online.

## ACKNOWLEDGEMENT

This work utilized the computational resources of the NIH HPC Biowulf cluster. (http://hpc.nih.gov).

