## Supplementary Material for "Coarse-Grained Modelling of DNA Plectoneme Pinning in the Presence of Base-Pair Mismatches"

### Coarse-Grained Modeling of DNA Plectoneme in the Presence of Base-Pair Mismatches: Supporting Information

**S1. Comparison of DNA bending energy at a G:T mismatch obtained with the OxDNA and all-atom models.**

Our goal is to develop an approach within the oxDNA2 model to accurately model the buckling of DNA containing mismatched base pairs. To this end we first measure the free energy required to bend a 15 bp long DNA with different mismatches parameterized in oxDNA2 and compare the results to all atom simulations performed by Sharma et al.(1) We use the same sequence of DNA as used in Sharma et al(1). To calculate the free energy of bending we simulate a 15 bp long DNA with the mismatch located at the center ($8^{th}$ base pair). We employ umbrella sampling to efficiently sample the free energy landscape. The collective variable ($\theta$) is the angle defined between the center of mass of 3 blocks of 5 bp's (block 1: 2-5 bp, block 2: 6-10 bp, block 3: 11-14 bp), the mismatched base pair is located at the center of the DNA (base 8/23). The ends of the DNA are unconstrained. We then apply a harmonic constraint, of type

$$U(\theta) = \frac{k}{2}(\theta - \theta_{ref})^2$$

using the COLVARS(2) module in LAMMPS. Here $k = 0.064(\epsilon_{LJ}/rad^2)$. We carry out both forward and backward sampling of the free energy landscape by first gradually decreasing $\theta_{ref}$ from 180° to 90° in steps of 5°, and then increasing $\theta_{ref}$ from 90° to 180° in steps of 5°. We simulate each step for $10^4\tau_{LJ}$ (30.3 ns) of which the first $10^2\tau_{LJ}$(0.3 ns) are equilibration steps and not considered in subsequent data analysis. The MD simulations are carried out in the NVT ensemble using the Langevin thermostat with a damping factor of 0.5 $\tau_{LJ}$and 10 $\tau_{LJ}$ for the transitional and rotational degrees of freedom, respectively. The equations of motion are integrated using the velocity-Verlet algorithm with a timestep of $0.001\tau_{LJ}$ (3.03 fs). We can then extract the free energy associated with bending using the weighted histogram analysis method (WHAM)(3) as implemented by Grossfield(4).

Sharma et al.[1] analyzed the results of their all atom simulations using the definition of angle provided by the COLVARS module and the definition of angle used in the 3DNA software. Sharma et al.[1] also noted that their results are slightly affected by the definition of bend angle.

We use the bending free energy obtained by Sharma et al. using the 3DNA software to compare the oxDNA2 model with all atom simulation. The 3DNA software fits a liner helical axis to the two fragments of a bent DNA molecule. The bend angle is defined as the angle between these two helical axes. This definition of the bend angle closely resembles the definition of bend angle for a coarse-grained model like oxDNA2.

Fig. S1 shows that, for a G:T mismatch, the OxDNA2 model reproduces the reduction in local bending rigidity observed in an all atom simulation of DNA.

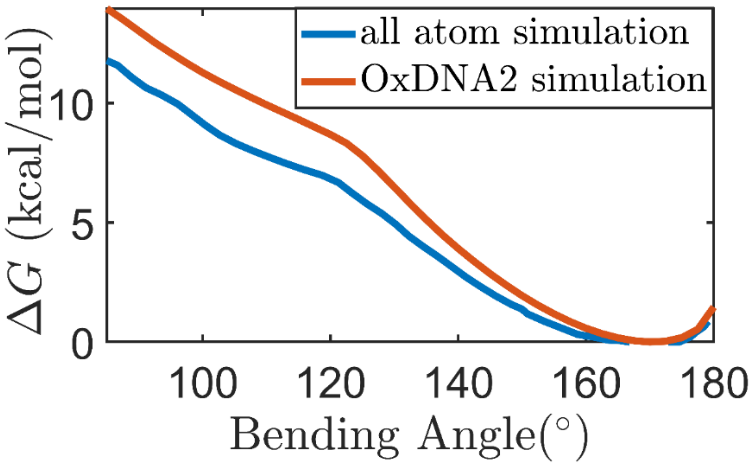


**Figure S1.** Comparison of free energy versus bend angle for a G:T mismatch obtained using an all atom model(1) (blue) and the OxDNA2 model (red).

**S2. Simulation of DNA under tension and no torsional stress**

We conduct simulations of stretched and untwisted DNA to calculate the twist stored in a DNA molecule with mismatched base pairs. We perform simulations for a 610 bp and 1010 bp DNA with either no mismatched bases, or with 2, 4, or 6 consecutive mismatched bases in the center of the DNA. We perform a single simulation for each mismatch size under different force and salt conditions. The procedure used to perform these simulations is similar to the procedure used for DNA under torsional stress. Each simulation for the 610 bp DNA under the high force-high salt condition is carried out for $2.3 \times 10^7 \tau_{LJ}$ $(69.69\ \mu s)$. Each simulation for the 610 bp DNA under the low force-low salt condition is carried out for $1.8 \times 10^7 \tau_{LJ}$ $(54.54\ \mu s)$, and simulations for 1010bp DNA under low force-low salt condition are carried out for $1.6 \times 10^7 \tau_{LJ}$ $(48.48\ \mu s)$.

We find that for the 2-mismatch case the twist at the mismatch is smaller as compared to intact DNA, which is presumably due the effect of flanking base pairs (i.e. the intact base pairs adjacent to the DNA). The flanking base pairs will have a larger effect on the twist for the 2-mismtach case as compared to the 4 and 6-mismatch cases. We still observe a torsional softening at the mismatched bases for the 2-mismatch case. Hence, when torsional stress is applied to the DNA with 2 mismatches, we observe a slightly higher change in the twist at the mismatch compared to the change in twist at the intact base pairs.

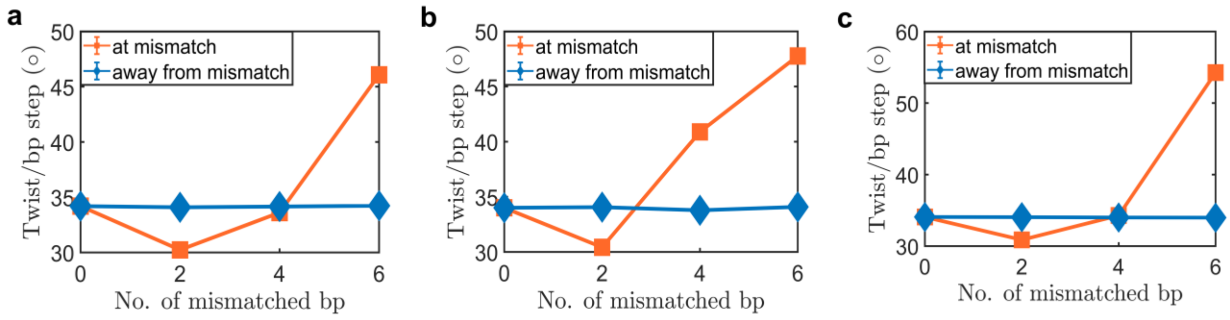


**Figure S2.** Average twist per bp step in degrees of a torsional stress free DNA as a function of the number of consecutive mismatched base pairs introduced at the DNA center for (a) 610 bp DNA with F=2 pN and 1 M monovalent salt, (b) 610 bp DNA with F=0.3 pN and 0.2 M monovalent salt, and (c) 1010 bp DNA with F=0.3 pN and 0.2M monovalent salt. Error bars (SEM) are smaller than the symbol size.

**S3. Mean squared displacement of plectoneme center**

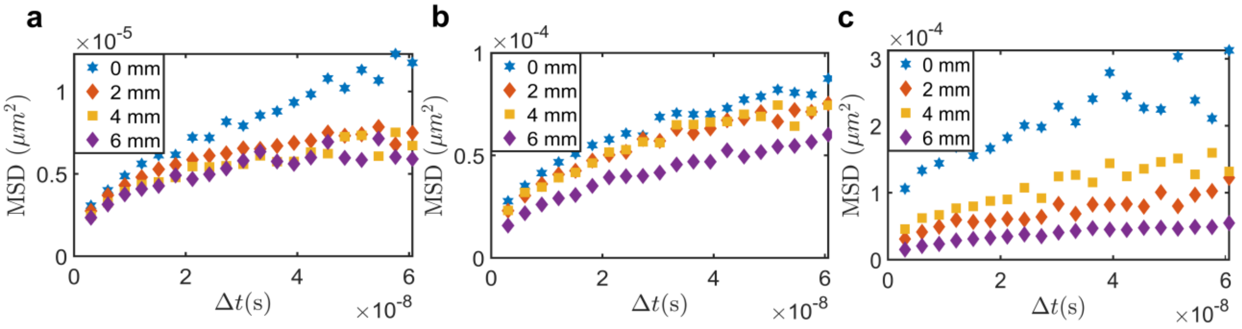


**Figure S3:** Mean squared displacement of the plectoneme center as a function of the number of consecutive mismatched base pairs introduced at the DNA center for (a) 610 bp DNA with F=2 pN and 1 M monovalent salt, (b) 610 bp DNA with F=0.3 pN and 0.2 M monovalent salt, and (c) 1010 bp DNA with F=0.3 pN and 0.2M monovalent salt. Note the change in the *y*-axis scaling. See Fig. 5 in the main text for examples of the plectoneme dynamics and the diffusion constants obtained by linear fitting the first 8 points of the MSD curves.

**S4. Comparison between simulation results and theoretical predictions based on theory developed by Brahmachari *et al*.**

Fig. S4 presents the probability of plectoneme pinning at the mismatch using the statistical-mechanical model developed by Brahmachari *et al.*(5). This model uses a structural parameter $\varepsilon$ to describe the size of the defect; $\varepsilon$ is defined as

$$\varepsilon = 1 - \frac{E_{\gamma^\dagger}}{E_\gamma} \qquad \text{(S1)}$$

Here, $E_{\gamma^\dagger}$ is the energy of a kinked end loop and $E_\gamma$ is the measure of energy of a defect free end loop. In order to compare the theoretical predictions with MD simulations, we calculate $\varepsilon$ using the results of the MD simulations. We use Fig. 2 b) to obtain the angle of the plectoneme end loop when the plectoneme is pinned at the mismatch and when it is not pinned at the mismatch. Fig. 1 (a) and Fig. 1 (b) can be used to obtain the energy required to bend a segment of DNA to a given angle. Using eq. S1 we can obtain an approximate estimate of $\varepsilon$ as a function of the number of mismatches in the simulated DNA molecule. We provide the values of the structural parameter $\varepsilon$ obtained from the simulations in table S1.

Note that the defect size parameter depends on the salt condition and the applied external force. The theoretical approach of Brahmachari *et al.* assumes the mismatches do not cause local torsional softening; however, we observe a slight torsional softening in the simulations. Although this torsional softening causes a small change in the critical linking number, it is unlikely to significantly affect the theoretical predictions. A second simplyfying assumption in the theoretical approach is considering the end loop fixed in size for a given force and defect size. In simulations, however, we observe variations in the end loop size. These fluctuations in the end loop may additionally destabilize the pinned plectoneme at lower forces.

An important parameter in the theoretical model for quantitative comparison with simulation results is the discretization length used to calculate discrete sliding states of the plectoneme. This discretization affects counting of states that are accessible via sliding of the plectoneme. A small discretization length leads to a large number of states resulting in a higher entropic cost to pinning of the plectoneme, and vice versa. This discretization length becomes significant in determining the stability of the pinned plectoneme state, especially for short DNA chains, the simulation case at hand. This characteristic length is expected to be much shorter than the persistence length of DNA, about 50 nm (the coarse-graining length scale for semiflexible polymer behavior), longer than the base pair length scale (about 1 nm), and comparable to other characteristic lengths in the system, such as the force-induced correlation length (about 10-20 nm); however, the exact numerical choice is not clear. We find that using a discretization length of 5-10 nm results in the best agreement with the simulations. The data displayed in Fig. S4 were obtained using 5 nm as the characteristic length distinguishing the sliding states of the plectoneme.

**Table S1. Structural parameters obtained from the simulation**

| | **N = 610 bp, F = 2 pN, S = 1 M & $\Delta \boldsymbol{Lk} = 3.46$** | **N = 610 bp, F = 0.3 pN, S = 0.2 M & $\Delta \boldsymbol{Lk} = 2.31$** | **N = 1010 bp, F = 0.3 pN, S = 0.2 M & $\Delta \boldsymbol{Lk} = 2.87$** |
|---|---|---|---|
| **Number of mismatches** | **$\boldsymbol{\varepsilon}$ obtained from simulations** | **$\boldsymbol{\varepsilon}$ obtained from simulations** | **$\boldsymbol{\varepsilon}$ obtained from simulations** |
| 0 | 0 | 0 | 0 |
| 2 | 0.23 | 0.06 | 0.1 |
| 4 | 0.5 | 0.26 | 0.32 |
| 6 | 0.75 | 0.41 | 0.55 |

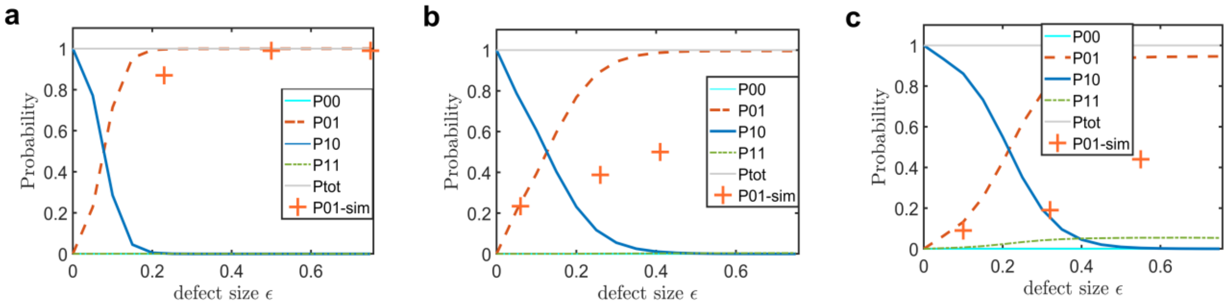


**Figure S4. Theoretical predictions of probability of plectoneme pining at the defect site.** Probability of the various plectoneme states: unbuckled (P00, cyan line), mobile (P10, blue line), pinned at the mismatch (P01, red dashed line), and the two-plectoneme state with one pinned and one mobile domain (P11, green dot-dashed line) are plotted as a function of the defect size. The orange crosses correspond to the probability of buckling at the mismatch obtained from MD simulations for defect sizes associated with 2, 4, and 6 mismatches (see table S1). (a) For the high force-high salt case with a 610 bp DNA, both theory and simulations indicate that 2, 4, or 6 mismatches stably pin the plectoneme. (b) For the low force-low salt case with a 610 bp long DNA, the theory predicts appreciable pinning only for 4 and 6 mismatches, whereas the 2 bp mismatch is unable to pin the plectoneme. The overall trend of increasing pinning probability with increasing mismatch size is recovered from simulations (Fig. 2), but there is an increasing discrepancy between theory and simulations for 4 and 6 mismatches. (c) For the low force-low salt case with a 1010 bp long DNA, the theory predicts appreciable pinning only for the 6 bp mismatch. Similar to the results with the 610 bp DNA under the low force and low salt conditions, the MD simulations agree with the theory for a 2 bp mismatch, but the probability of pinning the plectoneme at the mismatch increase more slowly with increasing mismatch size than predicted by theory.  P00 is zero for all values of $\varepsilon$ in all the three panels whereas P11 is zero for all values of $\varepsilon$ in panels (a) and (b).

#### S5. DNA sequences used for simulations

The specific sequences of DNA in the current work are provided below.

610 bp long DNA sequence:

AGAGTACTTAGGCTTGACGATTTCGCGCCTGAACTTCTGATAACTCAGTCTGAGAGACTAAGTTGACGTTCTATCCATCATCAGGT
GGGCTCAGAGATTGTGCGGCAGACTTAAGTGTAGTACCAGCTGCTGGTCAATTTGATCTATGCTGATCCGCTCGGAACGGGCCGT
GAAAGAAGTACTCTCGCCTATAGAACGGTTAGTGCTACGACTTTTGCGCGACACAATGTGGTAGTTATCTTCTGTTTTCCTGAATA
GTGAGCCTACCAGAAGAGGCCACCGACAAATCTGATGAGATAGATTTGGGATGTGTTTGCGGAGCCTCTGAAACGCTTGTTTATG
AGCAAGAGAGGTGCGGTGGGTATGACCGCCGTAGAAGTACCGTATTCTTCCGGGCTCGGTGGCAATGAACACTTAAGGGGCCGA
CACATTCTGAAGTCAATCGATGGACGGACCTCAACCGTGCACCCTTCTATATACGTGTGGCTAGGATACTCTAGCGTTTACCCGCC
GTCTTCCACGATGCCGAATATAAGCCGAGGATAAAGGTGCAGACAAATATCAGGCTTCGCAGTTGTGTAACTTCCTGTATTGTTGT
GCAGAGTACTTAAGAGTACTTAGGCTTGACGATTTCGCGCCTGAACTTCTGATAACTCAGTCTGAGAGACTAAGTTGACGTTCTAT
CCATCATCAGGTGGGCTCAGAGATTGTGCGGCAGACTTAAGTGTAGTACCAGCTGCTGGTCAATTTGATCTATGCTGATCCGCTCG
GAACGGGCCGTGAAAGAAGTACTCTCGCCTATAGAACGGTTAGTGCTACGACTTTTGCGCGACACAATGTGGTAGTTATCTTCTGT
TTTCCTGAATAGTGAGCCTACCAGAAGAGGCCACCGACAAATCTGATGAGATAGATTTGGGATGTGTTTGCGGAGCCTCTGAAAC
GCTTGTTTATGAGCAAGAGAGGTGCGGTGGGTATGACCGCCGTAGAAGTACCGTATTCTTCCGGGCTCGGTGGCAATGAACACTT
AAGGGGCCGACACATTCTGAAGTCAATCGATGGACGGACCTCAACCGTGCACCCTTCTATATACGTGTGGCTAGGATACTCTAGC
GTTTACCCGCCGTCTTCCACGATGCCGAATATAAGCCGAGGATAAAGGTGCAGACAAATATCAGGCTTCGCAGTTGTGTAACTTC
CTGTATTGTTGTGCAGAGTACTTA

1010 bp long DNA sequence:

AGAGTACTTAGGCTTGACGATTTCGCGCCTGAACTTCTGATAACTCAGTCTGAGAGACTAAGTTGACGTTCTATCCATCATCAGGT
GGGCTCAGAGATTGTGCGGCAGACTTAAGTGTAGTACCAGCTGCTGGTCAATTTGATCTATGCTGATCCGCTCGGAACGGGCCGT
GAAAGAAGTACTCTCGCCTATAGAACGGTTAGTGCTACGACTTTTGCGCGACACAATGTGGTAGTTATCTTCTGTTTTCCTGAATA
GTGAGCCTACCAGAAGAGGCCACCGACAAATCTGATGAGATAGACGGGAACACGGTTTGCGGAGCCTCTGAAACGCTTGTTTATG
AGCAAGAGAGGTGCGGTGGGTATGACCGCCGTAGAAGTACCGTATTCTTCCGGGCTCGGTGGCAATGAACACTTAAGGGGCCGA
CACATTCTGAAGTCAATCGATGGACGGACCTCAACCGTGCACCCTTCTATATACGTGTGGCTAGGATACTCTAGCGTTTACCCGCC
GTCTTCCACGATGCCGAATATAAGCCGAGGATAAAGGTGCAGACAAATATCAGGCTTCGCAGTTGTGTAACTTCCTGTATTGTTGT
GCAGAGTACTTAGGCTTGACGATTTCGCGCCTGAACTTCTGATAACTCAGTCTGAGAGACTAAGTTGACGTTCTATCCATCATCAG
GTGGGCTCAGAGATTGTGCGGCAGACTTAAGTGTAGTACCAGCTGCTGGTCAATTTGATCTATGCTGATCCGCTCGGAACGGGCC
GTGAAAGAAGTACTCTCGCCTATAGAACGGTTAGTGCTACGACTTTTGCGCGACACAATGTGGTAGTTATCTTCTGTTTTCCTGAA
TAGTGAGCCTACCAGAAGAGGCCACCGACAAATCTGATGAGATAGACGGGAACACGGTTTGCGGAGCCTCTGAAACGCTTGTTTA
TGAGCAAGAGAGGTGCGGTGGGTATGACCGCCGTAGAAGTACCGTATTCTTCCGGGCTCGGTGGATGCGAGAGTACTTAGGCTTG

ACGATTTCGCGCCTGAACTTCTGATAACTCAGTCTGAGAGACTAAGTTGACGTTCTATCCATCATCAGGTGGGCTCAGAGATTGTG
CGGCAGACTTAAGTGTAGTACCAGCTGCTGGTCAATTTGATCTATGCTGATCCGCTCGGAACGGGCCGTGAAAGAAGTACTCTCG
CCTATAGAACGGTTAGTGCTACGACTTTTGCGCGACACAATGTGGTAGTTATCTTCTGTTTTCCTGAATAGTGAGCCTACCAGAAG
AGGCCACCGACAAATCTGATGAGATAGACGGGAACACGGTTTGCGGAGCCTCTGAAACGCTTGTTTATGAGCAAGAGAGGTGCG
GTGGGTATGACCGCCGTAGAAGTACCGTATTCTTCCGGGCTCGGTGGCAATGAACACTTAAGGGGCCGACACATTCTGAAGTCAA
TCGATGGACGGACCTCAACCGTGCACCCTTCTATATACGTGTGGCTAGGATACTCTAGCGTTTACCCGCCGTCTTCCACGATGCCG
AATATAAGCCGAGGATAAAGGTGCAGACAAATATCAGGCTTCGCAGTTGTGTAACTTCCTGTATTGTTGTGCAGAGTACTTAGGC
TTGACGATTTCGCGCCTGAACTTCTGATAACTCAGTCTGAGAGACTAAGTTGACGTTCTATCCATCATCAGGTGGGCTCAGAGATT
GTGCGGCAGACTTAAGTGTAGTACCAGCTGCTGGTCAATTTGATCTATGCTGATCCGCTCGGAACGGGCCGTGAAAGAAGTACTC
TCGCCTATAGAACGGTTAGTGCTACGACTTTTGCGCGACACAATGTGGTAGTTATCTTCTGTTTTCCTGAATAGTGAGCCTACCAG
AAGAGGCCACCGACAAATCTGATGAGATAGACGGGAACACGGTTTGCGGAGCCTCTGAAACGCTTGTTTATGAGCAAGAGAGGT
GCGGTGGGTATGACCGCCGTAGAAGTACCGTATTCTTCCGGGCTCGGTGGATGCGCC

**S6. Sequence effect on DNA pinning**

Kim et al.(6) have shown that DNA sequence can influence plectoneme pinning. They found that intrinsic bend of a DNA sequence can be a significant factor in determining plectoneme pinning. We study the intrinsic bend in the DNA sequence used, to ensure we do not have a significantly bent section in the simulated DNA.

Previous studies (6) used dinucleotide steps to calculate the intrinsic bend in a DNA segment. Here instead of using the dinucleotide step method, we explicitly calculate the bend angle at various bp steps. The dinucleotide step method does not account for force applied to DNA and how that might affect the intrinsic bend at various base-pairs. To overcome these limitations, we explicitly calculated the intrinsic bend at various base pairs.

To calculate the intrinsic bend, we use the trajectories obtained from simulations of DNA under no torsional stress (same simulations used for results provided in supplementary section S2). To calculate bend at a bp, we use the same method used to calculate the end loop angle as described in the Methods section of the main text. Briefly, the bend angle at a base-pair is defined between the center of mass of 3 blocks of 5 bp. If the bp is at position *i*, the 3 blocks would be block 1: *i*-12 *to* *i*-8 bp, block 2: *i*-2 *to* *i*+2 bp, block 3: *i*+8 *to* *i*+12 bp.

Fig. S5 provides the bend angles at each base pair along the DNA. We find for intact DNA (0 mismatches) the bend angle is almost constant along the DNA.

For a DNA with mismatches, we find that the bend angle is significantly lower (i.e the DNA is highly bent) at the location of mismatch. We also find that the bend angle at the mismatch decreases with more mismatches.

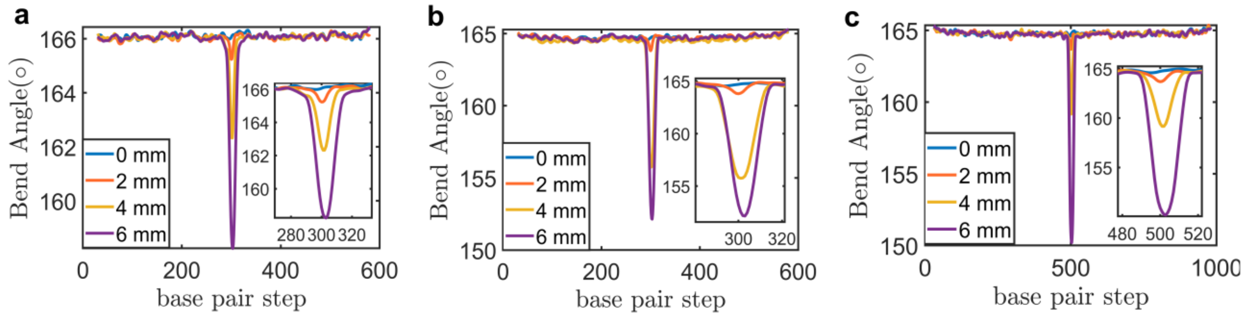


**Figure S5. Intrinsic bend. (a-c)** Average angle at each bp along the DNA backbone for different number of mismatches introduced at the center of the DNA for (a) 610 bp DNA with F=2 pN and 1 M monovalent salt, (b) 610 bp DNA with F=0.3 pN and 0.2 M monovalent salt, and (c) 1010 bp DNA with F=0.3 pN and 0.2 M monovalent salt. Insets provide a magnified view of the angle at the location of mismatches.
